# Building, Benchmarking, and Exploring Perturbative Maps of Transcriptional and Morphological Data

**DOI:** 10.1101/2022.12.09.519400

**Authors:** Safiye Celik, Jan-Christian Hütter, Sandra Melo Carlos, Nathan H Lazar, Rahul Mohan, Conor Tillinghast, Tommaso Biancalani, Marta M Fay, Berton A Earnshaw, Imran S Haque

## Abstract

1

The continued scaling of genetic perturbation technologies combined with high-dimensional assays such as cellular microscopy and RNA-sequencing has enabled genome-scale reverse-genetics experiments that go beyond single-endpoint measurements of growth or lethality. Datasets emerging from these experiments can be combined to construct perturbative “maps of biology”, in which readouts from various manipulations (e.g., CRISPR-Cas9 knockout, CRISPRi knockdown, compound treatment) are placed in unified, relatable embedding spaces allowing for the generation of genome-scale sets of pairwise comparisons. These maps of biology capture known biological relationships and uncover new associations which can be used for downstream discovery tasks. Construction of these maps involves many technical choices in both experimental and computational protocols, motivating the design of benchmark procedures to evaluate map quality in a systematic, unbiased manner. Here, we (1) establish a standardized terminology for the steps involved in perturbative map building, (2) introduce key classes of benchmarks to assess the quality of such maps, (3) construct maps from four genome-scale datasets employing different cell types, perturbation technologies, and data readout modalities, (4) generate benchmark metrics for the constructed maps and investigate the reasons for performance variations, and (5) demonstrate utility of these maps to discover new biology by suggesting roles for two largely uncharacterized genes.

**Author Summary:** With the proliferation of genetic perturbation, laboratory robotics, computer vision and sequencing technologies, a growing number of researchers are producing datasets that capture digital readouts of cellular responses to genetic perturbations at the full-genome-scale. Since each of these efforts utilizes different cellular models, experimental approaches, terminology, code bases, analysis methods and quality metrics, it is exceptionally difficult to reason through the pros and cons of possible design choices or even discuss the primary considerations when embarking on such an endeavor. These datasets can be powerful discovery tools to look at known biological relationships and uncover new associations in an unbiased manner, but only when paired with a computational pipeline to assemble the data into a digestible format. Moreover, there is great promise in looking across these data to highlight commonalities and differences that may be attributed to experimental or analytical approaches or the biological context. Therefore, a unified framework is necessary to align this nascent field and speed progress in assessing technologies and methods.

In this work we define a unified framework for building and benchmarking these perturbative maps, benchmark four different datasets assembled into 18 different maps, explore the impact of different design decisions and demonstrate how these maps can be used to elucidate gene functions. The framework we propose highlights the necessary steps for building any such map - embedding, filtering, aligning, aggregating and relating the data across perturbations. For benchmarking, we propose two main types of metrics and give examples which highlight the impact of different processing pipelines. Finally, we explore these maps to demonstrate their utility for confirming known biological relationships and nominating annotations for genes with unknown function.

We expect that this work will positively impact the nascent field of perturbative map building by enabling easier comparisons within and between technologies and methods through a shared language. Additionally, the associated code base is openly available and flexible enough to be easily extended with new methods, so we hope that it will become a resource for future researchers working on developing both laboratory and computational methodology. While there are too many confounding variables to make recommendations on the strengths of different technologies and cellular models at this time, highlighting that fact may prompt studies designed with the goal of directly comparing methods while holding other confounding variables fixed. Moreover, as the number of perturbative maps grows, the field will naturally consider the advantages of combining maps across modalities and the framework provided here can also help guide the evaluation of those efforts.

## 3 Introduction

Advances in genome editing technologies and high-throughput screening capabilities have enabled building perturbative maps through unbiased, large-scale profiling of genetic perturbations. These maps have massive potential to uncover novel biology and, when paired with compound screening, to accelerate drug discovery processes. The increasing prevalence of data generated by genetic perturbation technologies combined with high-dimensional assays [1, 2, 3, 4, 5, 6] signals a wider recognition of the power of these data to untangle complex biological mechanisms and to spur advancements in drug discovery.

Several recent studies have utilized single-cell pooled screening techniques to create genome-scale perturbation datasets. Replogle et al. [1] used pooled CRISPR interference (CRISPRi) libraries to generate genome-wide perturbation data with single cell RNA-seq as the readout. Ramezani et al. [2], Sivanandan et al. [3] and Funk et al. [4] used pooled CRISPR-Cas9-mediated gene knockouts with cellular imaging [7] followed by in-situ sequencing of the molecular barcode to assign guides to individual cells and obtain single-cell phenotypic perturbation readouts. While pooled screens are typically resource-efficient and cost-effective, it is not possible to tightly control the number of cells transfected with each specific guide RNA and to account for the interactions between cells that receive different perturbations. In contrast, recent studies [5, 6] utilized array-based screening where highly-automated labs apply thousands of distinct perturbations to separate cell populations in multi-well plates using CRISPR-Cas9 constructs that target individual genes.

These studies each generated map-building pipelines and analyses for their proposed datasets. However, to our best knowledge, no study has yet explored different genome-wide perturbative maps in a comprehensive manner using a shared language and consistent benchmarking measures, allowing for a comparison of different maps, particularly the computational pipeline choices for a given perturbation dataset.

Here, we first establish a systematic framework and vocabulary (Figure 1), as well as a codebase (github.com/recursionpharma/EFAAR_benchmarking), for constructing and evaluating such maps, which we expect will lead to more comparable analysis and optimization of maps going forward. We describe perturbation signal benchmarks for assessing the effect and consistency of individual perturbations, and biological relationship benchmarks for assessing biological relevance of a map through its ability to recapitulate annotated relationships from large-scale public sources. It is important to note that it is not our goal to establish the optimal analytical choices for each perturbation dataset, but to demonstrate how the benchmarks we present can to be used to optimize the analytical choices for each map.

**Figure 1:**
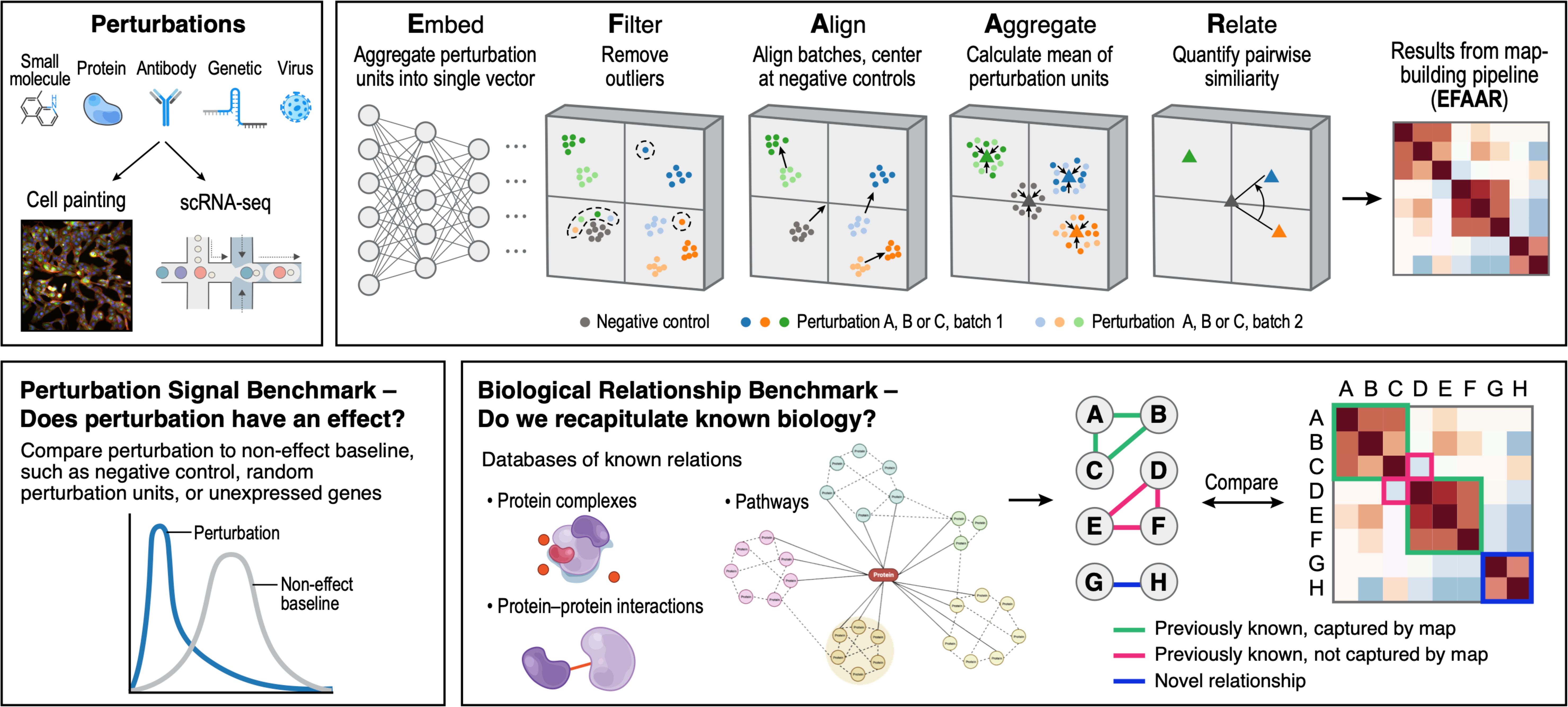
Graphical abstract of the introduced framework

We then construct maps from four recent gene perturbation datasets which perturb at least ∼8,000 genes each and have different perturbation modalities, readouts, and experimental setups. We then use our benchmarking framework to compute metrics for multiple computational analysis pipelines on each of these datasets. We use these metrics to explore the strengths and weaknesses of different pipelines in order to provide a deeper understanding of their performance and applicability. An important contribution of our study is the reporting of results from various annotation sources when examining the biological relationship benchmarks. While previous studies focused on recapitulation of protein complexes, we add in protein-protein interactions derived from pathways (Reactome) and signaling cascades (SIGNOR). With the maps we have constructed and analyzed, we provide compelling evidence for a relationship between *C18orf21*, an uncharacterized gene, and *ABT1*, *DKC1*, *POP1*, *POP5*, *RPP30*, and *UTP23*, suggesting that *C18orf21* participates in the RNase mitochondrial RNA processing complex. Similarly, we provide evidence for *C1orf131* interacting with *AATF*, *DDX52*, *NOL6*, *PDCD11*, *PNO1*, *RRP12*, and *UTP20*, supporting its involvement in the small subunit processome.

## 4 Results

### 4.1 Map building pipeline

Throughout this paper, we refer to a series of experiments involving genome-scale perturbations as a *perturbation **dataset*** and the final set of perturbation embeddings and associations between these as a *perturbative **map***. In the main results of this work, we present 18 maps constructed on four datasets.

We call the smallest experimental entity that is measured in a map context a *perturbation unit*. This is a cell in single-cell transcriptomic assays or in pooled screens where each cell receives a distinct barcoded perturbation [1, 8, 3, 2]. In the arrayed setting, a perturbation unit is a well containing hundreds of cells treated with the same perturbation [7, 9, 6]. Each perturbation unit is associated with assay data (e.g., images or gene transcript counts). Building a map which relates perturbations in a meaningful way from these raw assay data requires a number of post-experimental processing steps which we divide into five categories and refer to as an EFAAR pipeline.

- **E**mbedding assay data from each perturbation unit to generate a tractably-sized numeric representation
- **F**iltering perturbation units that do not pass quality criteria
- **A**ligning different batches of perturbation units
- **A**ggregating replicate units representing each targeted perturbation (e.g., a gene)
- **R**elating different perturbations to each other with one or more numeric values

#### 4.1.1 Embedding

This step is aimed at reducing high-dimensional assay data (e.g. 20,000 gene expression values or over one million image pixel values) to a tractable numerical representation that can be used for downstream tasks. For imaging data, embeddings may come from morphological features derived using software like CellProfiler [10] or from intermediate layers of neural networks[5]. For expression data, linear dimensionality reduction methods like principal component analysis (PCA) or non-linear methods based on neural networks may be used [11, 12].

#### 4.1.2 Filtering

In any experimental screening process, some perturbation units will not satisfy pre-defined quality criteria and need to be filtered out. This filtering can occur before or after embeddings are generated. Examples include wells with too high or too low pixel intensity or cells that receive more than one target guide.

#### 4.1.3 Aligning

A *batch effect* is a systematic bias shared by all observations obtained under similar experimental conditions (e.g., microscopy acquisition artifacts, cell donor batch, incubation times) that potentially confound the interpretation of desired biological signal. Aligning across batches can reduce the effects of these unintended variations and bring out more biologically-meaningful relationships.

A baseline approach for aligning perturbation units is to use control units in each batch to center and scale features. Another linear method aligning not only the first order statistics but also the covariance structures is Typical Variation Normalization (TVN) [13]. Non-linear methods based on nearest neighbor matching [14, 15] or variational autoencoders have been particularly successful for the alignment of single cell transcriptomic data [11, 12] as well as modeling batch effects in image data [16, 17, 18]. Another important technique for image data is Instance Normalization [19] which involves normalizing the features across each channel in an individual sample. This aids in removing bias in feature statistics when training data is divided into small computational batches. In gene expression domain, probabilistic batch effect correction methods that jointly model contributions from genotype and confounding factors have been used to map expression quantitative trait loci (eQTLs) [20]. ComBat and ComBat-seq [21, 22] utilize empirical Bayes models to adjust for batch effects and have been widely used for gene expression data.

#### 4.1.4 Aggregating

There are typically multiple technical or biological replicates representing each perturbation in a given dataset, e.g. the same perturbation may be applied to dozens of wells or hundreds of cells and these must be combined in order to produce a final representation of a perturbation. Coordinate-wise mean and median aggregation are commonly used, while more advanced methods like the Tukey median [23] may reduce the impact of outliers on the final representation, but increase computational complexity.

#### 4.1.5 Relating

Identifying relationships between biological entities (e.g., gene-gene interactions arising from protein complexes or signaling pathways) is an important use case for maps built based on genetic perturbations. Computing distances (e.g., Euclidean distance) or other similarity or dissimilarity measures (e.g. cosine similarity) between aggregated perturbation representations is commonly used as a proxy for relationships. These, in turn, can also be used to visualize the global structure of perturbations through further dimensionality reduction techniques such as uniform manifold approximation (UMAP) [24] or minimum-distortion embedding (MDE) [25].

These EFAAR steps may take place in a different order, multiple times (e.g., perturbation units may be filtered pre- and post-embedding), or potentially in a single end-to-end process. We highlight these steps as essential when constructing any perturbative map.

### 4.2 Map benchmarking pipeline

To assess maps built using different perturbation technologies, different readout data modalities, and different EFAAR pipelines, it is important to benchmark the resulting perturbation representations both in terms of the effect of the individual perturbations and interactions between perturbations. We call these “perturbation signal benchmarks” and “biological relationship benchmarks”, respectively.

#### 4.2.1 Perturbation signal benchmarks

Perturbation signal benchmarks assess the consistency and magnitude of the representations of individual perturbations in a map. In this work, consistency is measured by the average cosine of the angle between replicates of a perturbation (Section 6.6), and magnitude is measured by the energy distance [26, 27] between the control and perturbation samples (Section 6.7).

Our perturbation signal benchmarks identify the genes whose representation achieves a p-value < .05 in the associated statistical test using the unexpressed genes as the baseline of no signal. (Section 6.8). The fraction of such genes can be compared between different map processing pipelines (EFAAR parameter choices) and stratified by global annotations like gene expression or functional gene groups. Genes with a significant p-value for both consistency and magnitude receive higher priority when selecting hypotheses, such as identifying targets to pursue in downstream drug discovery tasks.

#### 4.2.2 Biological relationship benchmarks

A typical use case for a map of biology is to discover novel, biologically-relevant relationships. The following five annotation sources are used to assess the degree to which each map detects biological relationships. The underlying hypothesis is that if a map can identify known relationships to a high degree, it is an indication that it demonstrates a strong representation of existing biology, and is therefore more likely to accurately represent and uncover novel biological relationships.

- **CORUM** (COmprehensive ResoUrce of Mammalian): gene clusters representing protein complexes [28]
- **HuMAP**: gene clusters representing protein complexes [29]
- **Reactome**: protein-protein interactions derived from pathways [30]
- **SIGNOR** (SIGnaling Network Open Resource): signaling pathway interactions [31]
- **StringDB**: functional protein-protein associations [32]

Recall of annotated pairs is reported for the most extreme 10% of pairwise relationships (Section 6.9). The main premise is that genes whose protein products have related functions or that act in concert will produce readouts that, when properly processed, display geometric relationships (e.g., similarity). We consider 5% from both tails of the pairwise similarity distribution since negative relationships can indicate opposing functions between genes. A random, uninformative map would achieve a recall of 10%.

Across the five data sources we utilize for benchmarking, there are a total of 143,252 unique relationships. 29,600 of these relationships exist in at least two sources, and 54 of them exist in all five sources (Supplementary Figure 2).

In this study we analyze maps of gene perturbations, as multiple genome-scale datasets have recently become available ([1, 5, 6, 2]). The framework presented here can also be applied to datasets with other types of perturbations. For example, when a map includes both genes and small molecules at a large scale, sources annotating relationships between small molecules and their target genes ([33, 34]) can be used for benchmarking.

### 4.3 Applications of the map building and benchmarking framework

#### 4.3.1 Datasets

We built and benchmarked maps on four genome-scale CRISPR-based perturbation datasets to create transcriptional and morphological maps using the following data sets:

- **RxRx3** from Recursion contains deep neural network embeddings of phenomic images, where CRISPR-Cas9 was applied in an arrayed format to target ∼17,000 genes in primary HUVEC cells [5, 35].
- **GWPS** (Genome-Wide Perturb-Seq) contains single-cell RNA-seq counts as the readout, where CRISPRi was applied to knock down ∼10,000 expressed genes in K562 cells [1].
- **cpg0016** from the JUMP (Joint Undertaking of Morphological Profiling) consortium contains Cell Profiler features of Cell Painting [7] images, where CRISPR-Cas9 was applied in an arrayed format to target ∼8,000 genes in the U2OS cell line [6].
- **cpg0021**, contains Cell Profiler features of Cell Painting images, where CRISPR-Cas9 was applied in a pooled optical screening format to perturb ∼20,000 genes in HeLa cells [2].

Table 1 presents an overview of the properties of the four datasets. For the three morphology datasets (RxRx3, cpg0016, and cpg0021), details about cell painting protocols are in Supplementary Table 1.

**Table 1:**
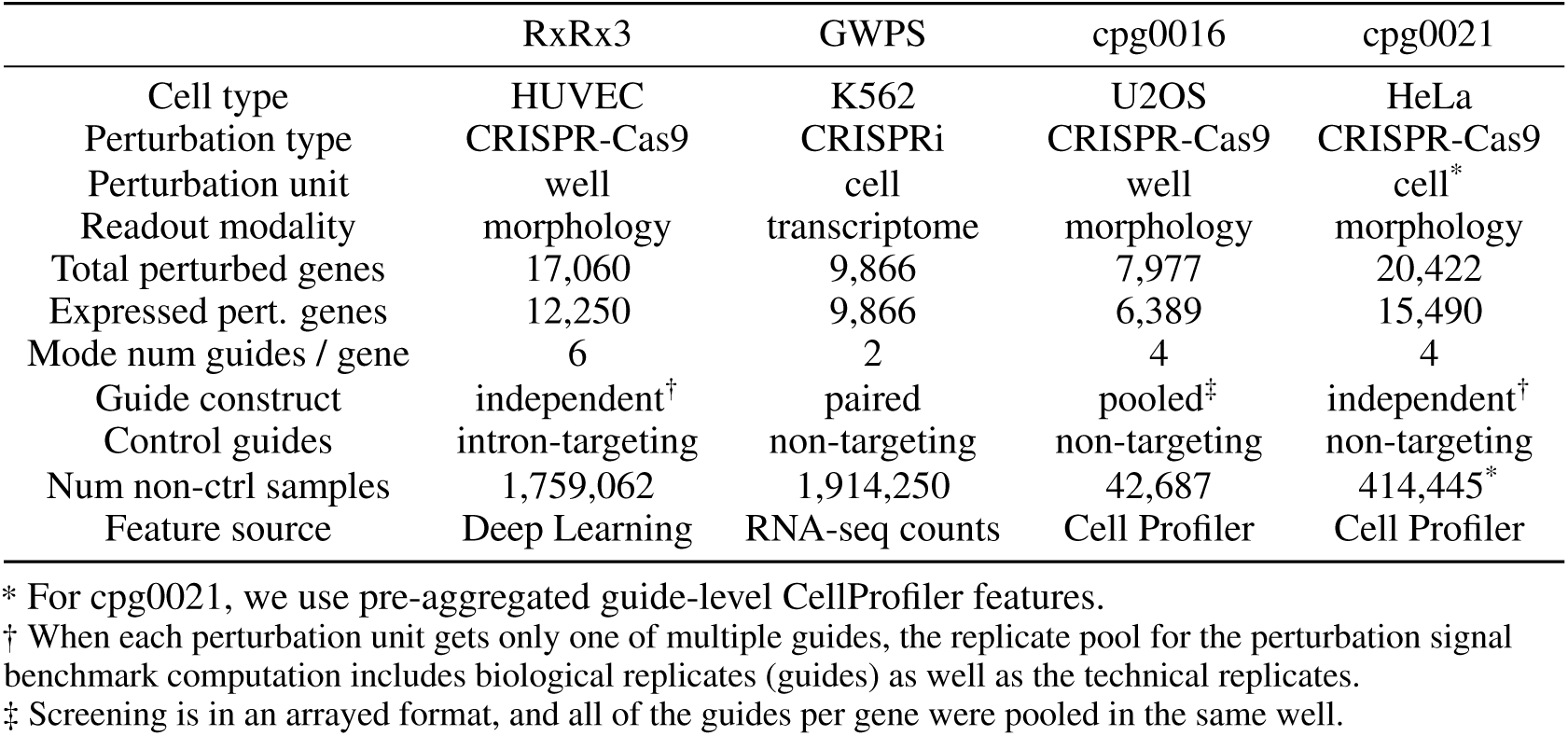
Properties of the four datasets on which we are applying the map building and benchmarking framework. These statistics specifically pertain to the portions of these datasets utilized in our application (Section 6.1) and not necessarily the full data available from each source. We use original study authors’ definition of “expressed” gene whenever applicable (Section 6.2).

|  | RxRx3 | GWPS | cpg0016 | cpg0021 |
| --- | --- | --- | --- | --- |
| Cell type | HUVEC | K562 | U2OS | HeLa |
| Perturbation type | CRISPR-Cas9 | CRISPRi | CRISPR-Cas9 | CRISPR-Cas9 |
| Perturbation unit | well | cell | well | cell* |
| Readout modality | morphology | transcriptome | morphology | morphology |
| Total perturbed genes | 17,060 | 9,866 | 7,977 | 20,422 |
| Expressed pert. genes | 12,250 | 9,866 | 6,389 | 15,490 |
| Mode num guides / gene | 6 | 2 | 4 | 4 |
| Guide construct | independent† | paired | pooled‡ | independent† |
| Control guides | intron-targeting | non-targeting | non-targeting | non-targeting |
| Num non-ctrl samples | 1,759,062 | 1,914,250 | 42,687 | 414,445* |
| Feature source | Deep Learning | RNA-seq counts | Cell Profiler | Cell Profiler |

\* For cpg0021, we use pre-aggregated guide-level CellProfiler features.
† When each perturbation unit gets only one of multiple guides, the replicate pool for the perturbation signal benchmark computation includes biological replicates (guides) as well as the technical replicates.
‡ Screening is in an arrayed format, and all of the guides per gene were pooled in the same well.

#### 4.3.2 EFAAR pipelines

Input data for the EFAAR pipelines were collected through the original sources of the four datasets (Section 6.1). The embedding and alignment steps for each dataset and processing pipeline are described below. Aggregation is performed by taking the mean of the aligned embeddings. Relationships are computed using the cosine similarity between the aggregated embeddings.

##### RxRx3 pipelines

- <u>Raw</u>: This refers to the 128-dimensional well-level RxRx3 embeddings. They were generated by passing images through a weakly-supervised convolutional neural network (CNN) pre-trained on a large set of proprietary Recursion data [36].
- <u>Raw-CS</u>: Raw RxRx3 embeddings are aligned via centering and scaling each feature to the mean and standard deviation of the controls in each batch.
- <u>Raw-TVN</u>: TVN [13] is applied on the RxRx3 embeddings after first centering and scaling features to the mean and standard deviation of the controls globally (not by batch). TVN aligns the data by PCA, centering and scaling globally, and correlation alignment (CORAL) [37] for each batch, all by controls.
- <u>PCA</u>: PCA is applied on the raw embeddings after centering and scaling all features in each batch, and all 128 principal components (PCs) are retained as the embeddings.
- <u>PCA-CS</u>: PCA-transformed embeddings are aligned via centering and scaling each batch by controls.
- <u>PCA-TVN</u>: A similar pipeline to the Raw-TVN, but applied on the PCA-transformed embeddings as the input instead of the raw embeddings.

##### GWPS pipelines

- <u>scVI</u>: Raw single-cell expression values are embedded to 128 dimensions using scVI (single-cell Variational Inference) [12]. scVI is a conditional variational auto-encoder providing both embedding and alignment in a single network (Section 6.3).
- <u>scVI-CS</u>: scVI embeddings are aligned via centering and scaling each batch by controls.
- <u>scVI-TVN</u>: A similar pipeline to the Raw-TVN in RxRx3, but applied on the scVI embeddings as the input.
- <u>PCA</u>: PCA is applied on the raw RNA-seq counts and the top 128 PCs are retained as the embeddings.
- <u>PCA-CS</u>: The same pipeline as in RxRx3.
- <u>PCA-TVN</u>: The same pipeline as in RxRx3.

##### cpg0016 pipelines

- <u>PCA</u>: PCA is applied on the filtered CellProfiler features (Section 6.4) and the top 128 PCs are retained as the embeddings.
- <u>PCA-CS</u>: The same pipeline as in RxRx3 and GWPS.
- <u>PCA-TVN</u>: The same pipeline as in RxRx3 and GWPS.

##### cpg0021 pipelines

- <u>PCA</u>: PCA is applied on the CellProfiler features which are pre-aggregated across cells to the guide level, and the top 128 PCs are retained as the embeddings.
- <u>PCA-CS</u>: The same pipeline as in RxRx3, GWPS, and cpg0016.
- <u>PCA-TVN</u>: The same pipeline as in RxRx3, GWPS, and cpg0016.

Here, we report results from pipelines generating 128-dimensional embeddings. Supplementary Table 2 includes biological relationship benchmarks for larger embedding spaces. The EFAAR choices above were made to analyze the four perturbation datasets as consistently as possible. This exploration is not intended to be all-encompassing; other methods might yield better results for different perturbation datasets.

#### 4.3.3 Benchmarking results

Here we look at the perturbation signal and biological relationship benchmarks for different EFAAR pipelines. We particularly check how sensitive these benchmarks are to the changes in the EFAAR steps. This facilitates the selection of the most effective pipeline for discovering novel relationships.

We restrict the benchmarks to the perturbed genes that are expressed in the respective cell type (Section 6.2). While we attempted to minimize differences between datasets with consistent EFAAR choices, values are not directly comparable across datasets due to inherent differences in cell type, mode of genetic perturbation, sample size, and assay design. However, observed trends for different EFAAR pipelines within a dataset can be informative, revealing strengths and weaknesses for different computational methodologies.

Figure 2 provides the perturbation signal (left) and biological relationship benchmarks (right) for the RxRx3 maps we constructed. TVN alignment shows an increased performance for all metrics and both of the embedding methods (raw and PCA). We believe this is because TVN is more effective at reducing batch-to-batch variation.

**Figure 2:**
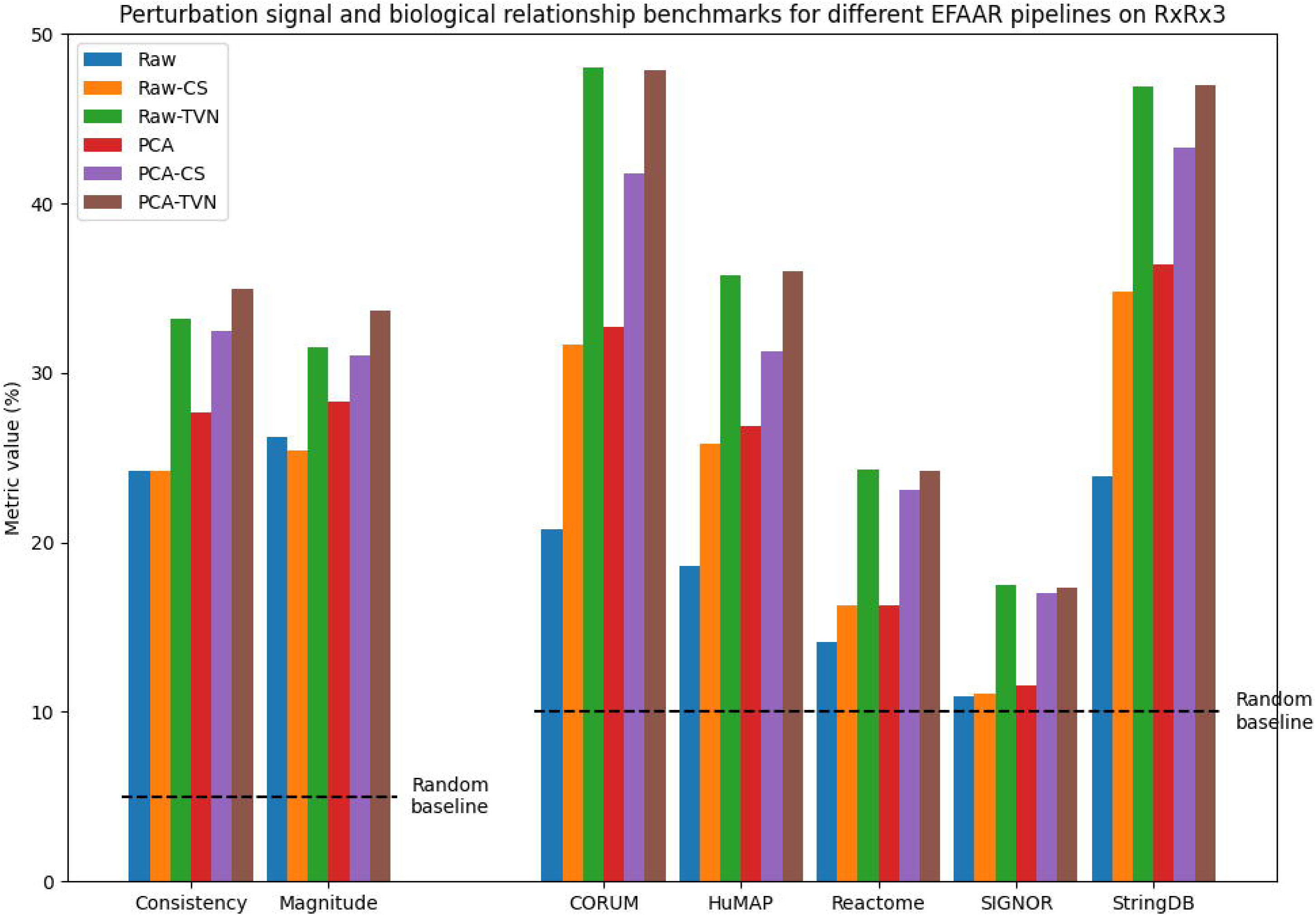
Benchmarking results in maps constructed using RxRx3 and different EFAAR pipelines. Bars for consistency and magnitude (left) show the percentage of perturbations with a significant p-value (< .05). CORUM, HuMAP, Reactome, SIGNOR and StringDB bars show the biological relationship benchmarks, i.e., the percentage of annotated relationships falling within the 5% tails (from each side) of the distribution of all pairwise cosine similarities. The random baseline for the biological relationship benchmarks is 10%.

Figure 3 shows benchmarks for different EFAAR pipelines on the GWPS dataset. Since this dataset does not include unexpressed gene perturbations, perturbation signal benchmarks cannot be calculated. Similar to the RxRx3 results earlier, using TVN for the alignment step leads to an increased performance for both of the embedding methods (scVI and PCA). Interestingly, the PCA embeddings show a better performance than scVI across all alignment methods.

**Figure 3:**
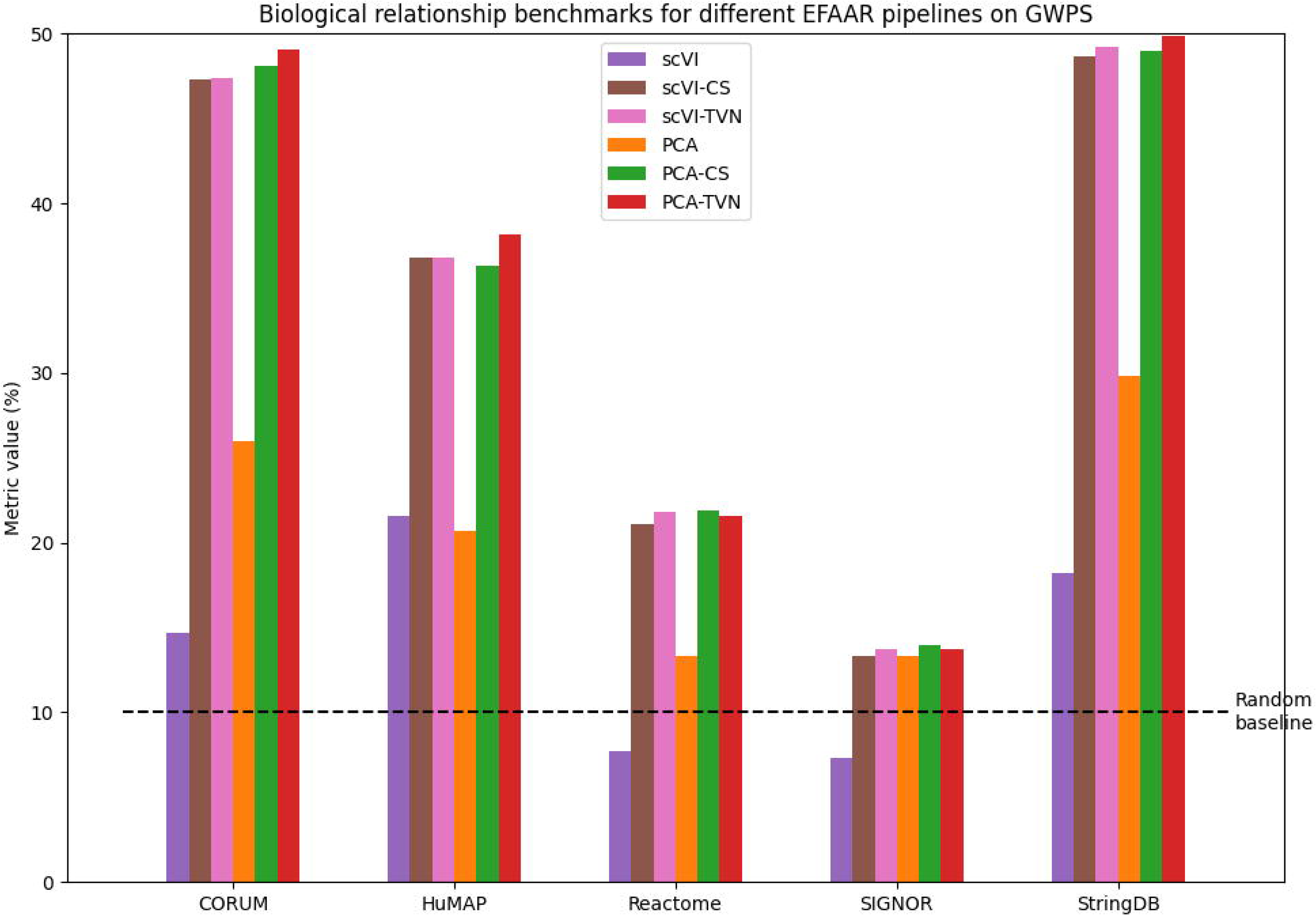
Biological relationship benchmarking results in maps constructed using different EFAAR pipelines on the GWPS dataset. Bar height shows the percentage of annotated relationships which fall within the 5% tails (from each side) of the distribution of all pairwise cosine similarities. The random baseline is 10%.

Figures 4 and 5 show the perturbation signal and biological relationship benchmark values for different EFAAR pipelines on the cpg0016 and cpg0021 datasets, respectively. Again, TVN alignment enhances the recapitulation of annotated gene-gene relationships for the majority of annotation sets (right), but PCA or PCA-CS performs better on perturbation signal benchmarks (left). For cpg0016, an improved batch alignment may have negative effects on perturbation signal benchmarks because the plate layouts are not fully randomized and plate position effects may add to the observed phenotype.

**Figure 4:**
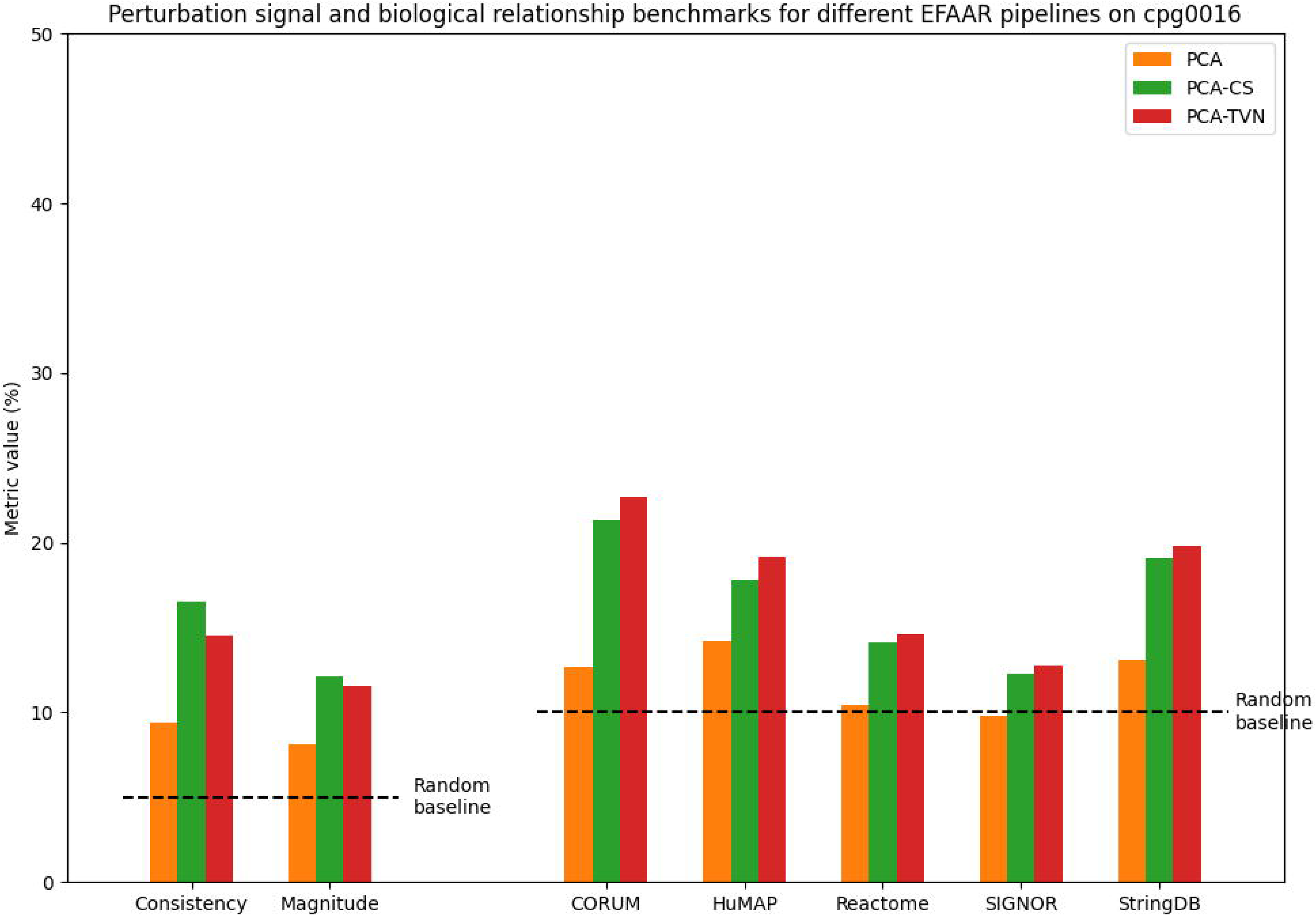
Benchmarking results in maps constructed using different EFAAR pipelines on the cpg0016 dataset. Bars for consistency and magnitude (left) show the percentage of perturbations with a significant p-value (< .05) for each measure. CORUM, HuMAP, Reactome, SIGNOR and StringDB bars show the biological relationship benchmarks, i.e., the percentage of annotated relationships falling within the 5% tails (from each side) of the distribution of all pairwise cosine similarities. The random baseline for the biological relationship benchmarks is 10%.

**Figure 5:**
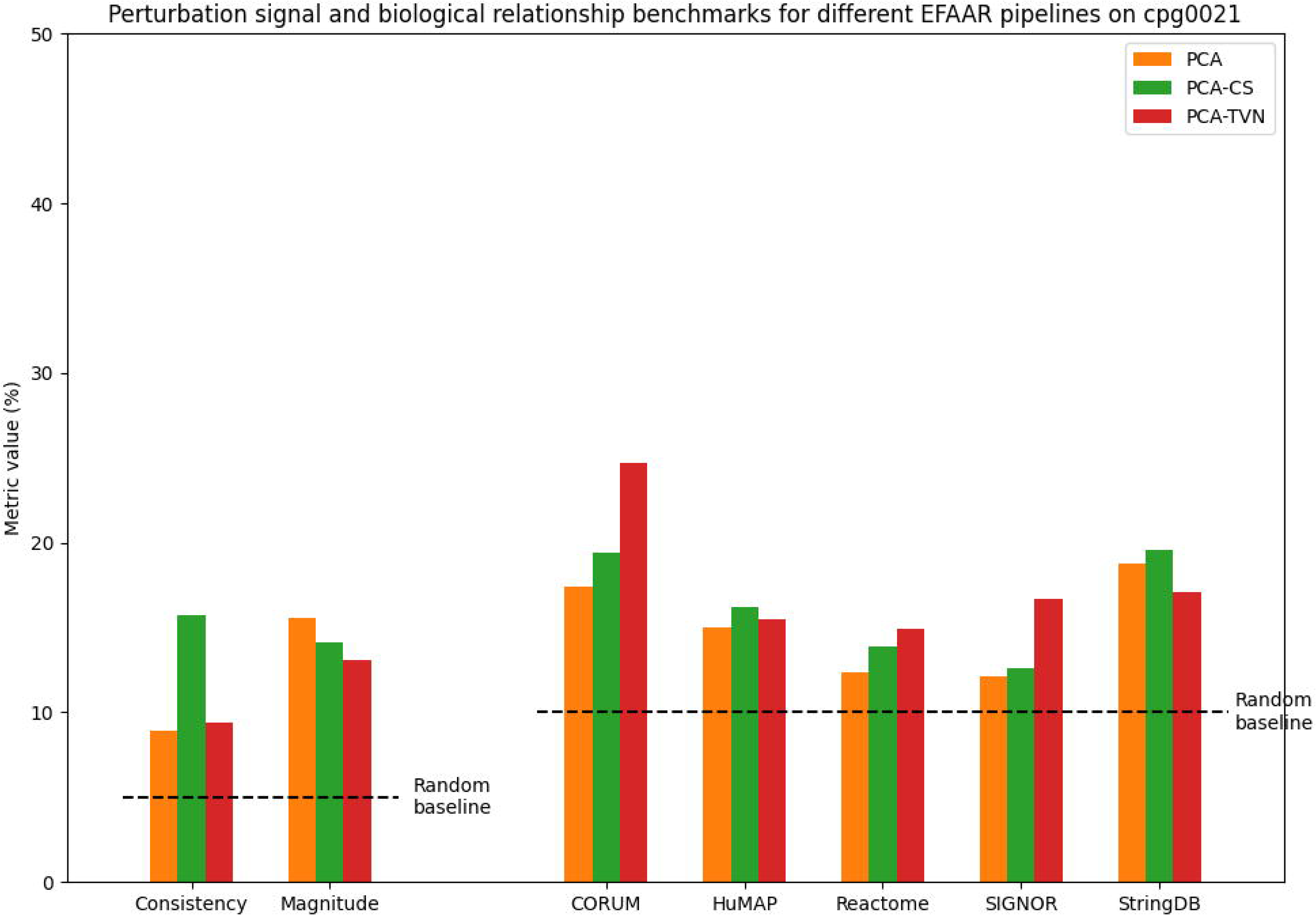
Benchmarking results in maps constructed using different EFAAR pipelines on the cpg0021 dataset. Bars for consistency and magnitude (left) show the percentage of perturbations with a significant p-value (< .05) for each measure. CORUM, HuMAP, Reactome, SIGNOR and StringDB bars show the biological relationship benchmarks, i.e., the percentage of annotated relationships falling within the 5% tails (from each side) of the distribution of all pairwise cosine similarities. The random baseline for the biological relationship benchmarks is 10%.

Looking across datasets, benchmark values are lower in the best-performing cpg0016 and cpg0021 maps than in the best-performing RxRx3 and GWPS maps. This could be due to a large number of differences across the datasets (Table 1). For example, although RxRx3, cpg0016, and cpg0021 are all morphology datasets that use CRISPR-Cas9 gene editing, RxRx3 employs intron-targeting guides as the negative controls. This approach may help distinguish between the effects of Cas9-induced cutting and the biological signals resulting from the knockout of the target gene. Alternatively, the single-guide treatments in RxRx3 wells with ∼1500 cells may contain more information or offer more stable representations of the gene perturbations than the single cells from cpg0021 or pooled-guide wells in cpg0016.

We further investigate how sample size influences the efficacy of maps in recapitulating biological relationships by randomly sub-sampling replicates (Section 6.10) for the PCA, PCA-CS, and PCA-TVN maps for all datasets. Increasing the sample size typically leads to an increased recall of known biology and reduced error (measured across different random subsets) (Supplementary Figures 3, 4, 5, 6). RxRx3 and cpg0021 show the largest improvement with increased replicates which may be due to these datasets measuring guides separately and thus having a more diversity across replicates.

#### 4.3.4 Perturbative maps built using different technologies surface different biology

To broadly assess the utility of each dataset, we examined which CORUM protein complexes the maps are able to identify (Section 6.11). We applied a stringent p-value cutoff of .01 during our exploration of maps for assessing their utility (this section) and using them for identifying poorly characterized cell functions (next section). Examined complexes are not mutually exclusive and may overlap; they may also be subsets of each other.

Figure 6a shows the overlaps of significantly identified complexes across the three maps with fully unblinded metadata (GWPS, cpg0016, and cpg0021). We utilized maps built using the PCA-TVN pipeline as these perform best on the biological relationship benchmarks (Figures 3, 4, 5). GWPS identifies the most complexes as well as the most unique complexes. It also identifies notably more complexes than the other two maps. The largest overlap occurs with complexes shared across GWPS and cpg0016, but not identified by cpg0021.

**Figure 6:**
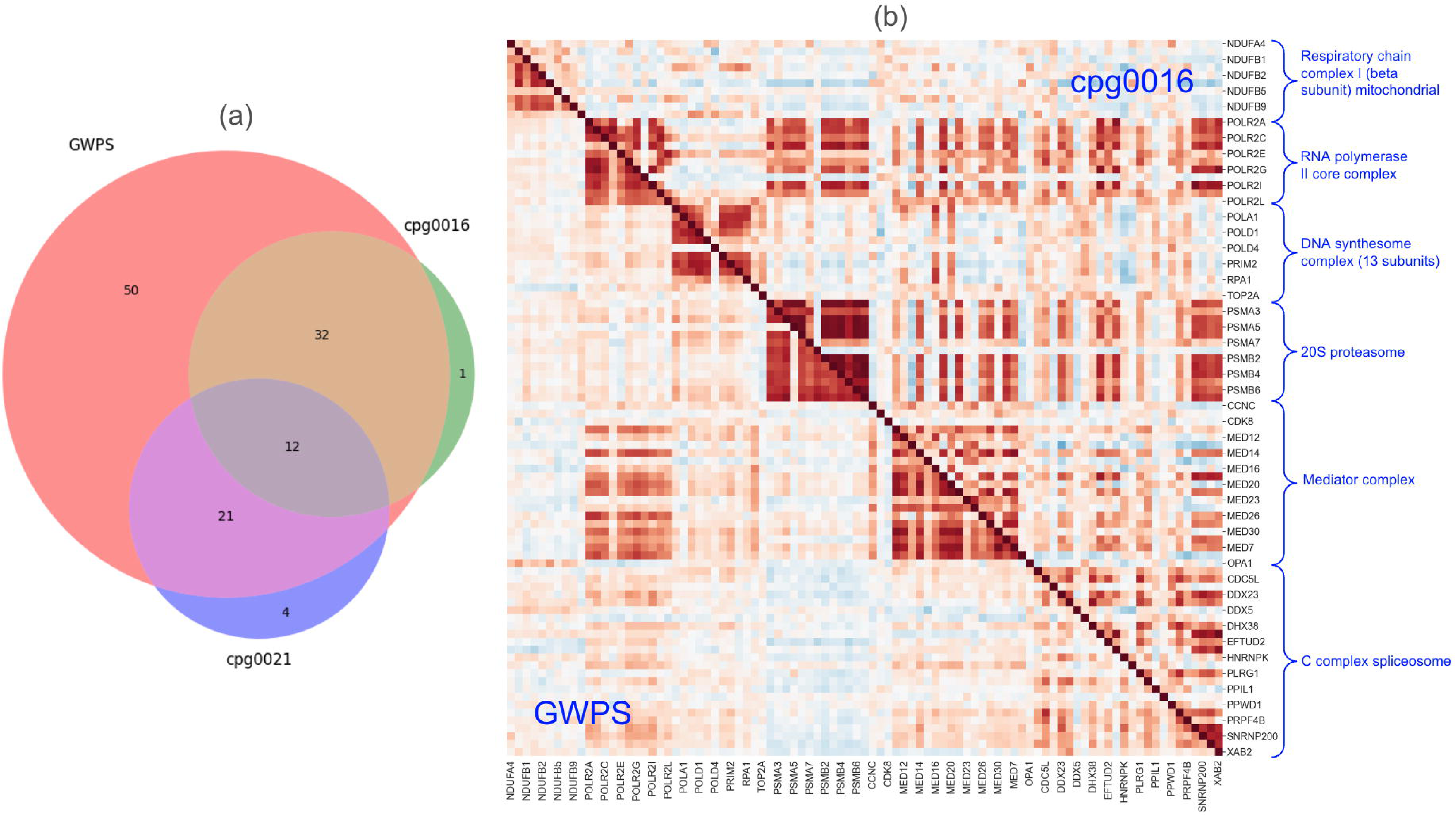
(a)Venn diagram of the intersection of the CORUM protein complexes captured by the PCA-TVN maps from each of GWPS, cpg0016, and cpg0021. There are 153 evaluated complexes (those with at least ten expressed genes) for GWPS, 83 for cpg0016, and 169 for cpg0021. (b) A split cosine similarity heatmap of the genes in six non-overlapping complexes out of the 12 identified by all of the GWPS, cpg0016, and cpg0021 maps. Below the diagonal represents similarities for the GWPS map, and above the diagonal represents similarities for the cpg0016 map.

Next, we examined complexes identified by all three maps, as well as those uniquely identified by one of the maps. 12 complexes are consistently identified by all three maps. These involve fundamental cellular processes crucial for the proper functioning and regulation of a cell and represent key molecular machinery involved in maintaining cellular integrity, survival, and functionality (Supplementary Table 3). To delve deeper, we generated a split heatmap of six of the 12 complexes in GWPS and cpg0016, the two maps identifying the most complexes. The 20S proteasome shows highly robust within-complex relationships in both transcriptomic and morphological maps, whereas the genes with RNA polymerase II core complex are similar to RNA polymerase II core complex and Mediator complex genes (Figure 6b). This data is consistent with the interaction between the Mediator complex and RNA polymerase II [38].

Fifty complexes are uniquely identified by the CRISPRi-based GWPS map with a transcriptional readout (Supplementary Table 4). A large number of these show associations with chromatin remodeling and transcriptional regulation. One complex is uniquely identified by the cpg0016 map, though the p-value for it is also near the significance level in the cpg0021 map (Supplementary Table 5). Three complexes are uniquely identified by the cpg0021 map (Supplementary Table 6), which is the largest of the three datasets with morphological readout (Table 1). The Emerin complex and Intraflagellar Transport Complex B have roles in maintaining cell shape and structure, thus directly impacting cell morphology.

Next, we examined the cosine similarity structure of the Integrator protein complex which was explored in the GWPS study [1] and is one of the fifty complexes uniquely identified by the GWPS map (Supplementary Table 4). The previously uncharacterized gene *C7orf26* was suggested to be a subunit of this complex by Drew et al. [29] and Replogle et al. [1], and was officially renamed *INTS15* in January 2022. We examined how well this complex’s structure is captured by the maps we constructed using other perturbation datasets. This complex was identified by RxRx3 (p-value: 3.6e-38) and by GWPS (p-value: 2.2e-25), but not by cpg0021 (p-value: .053). Integrator’s identification in the cpg0016 map cannot be assessed since only one of the genes in the complex was screened in the cpg0016 dataset. Figure 7 represents the cosine similarity heatmap for the Integrator complex in the RxRx3 and GWPS maps. Both of these maps accurately identify the modular structure of the Integrator complex and place C7orf26 in the enhancer module with *INTS10*, *INTS13*, and *INTS14*. Supplementary Figure 7 represents the heatmap for the cpg0021 map.

**Figure 7:**
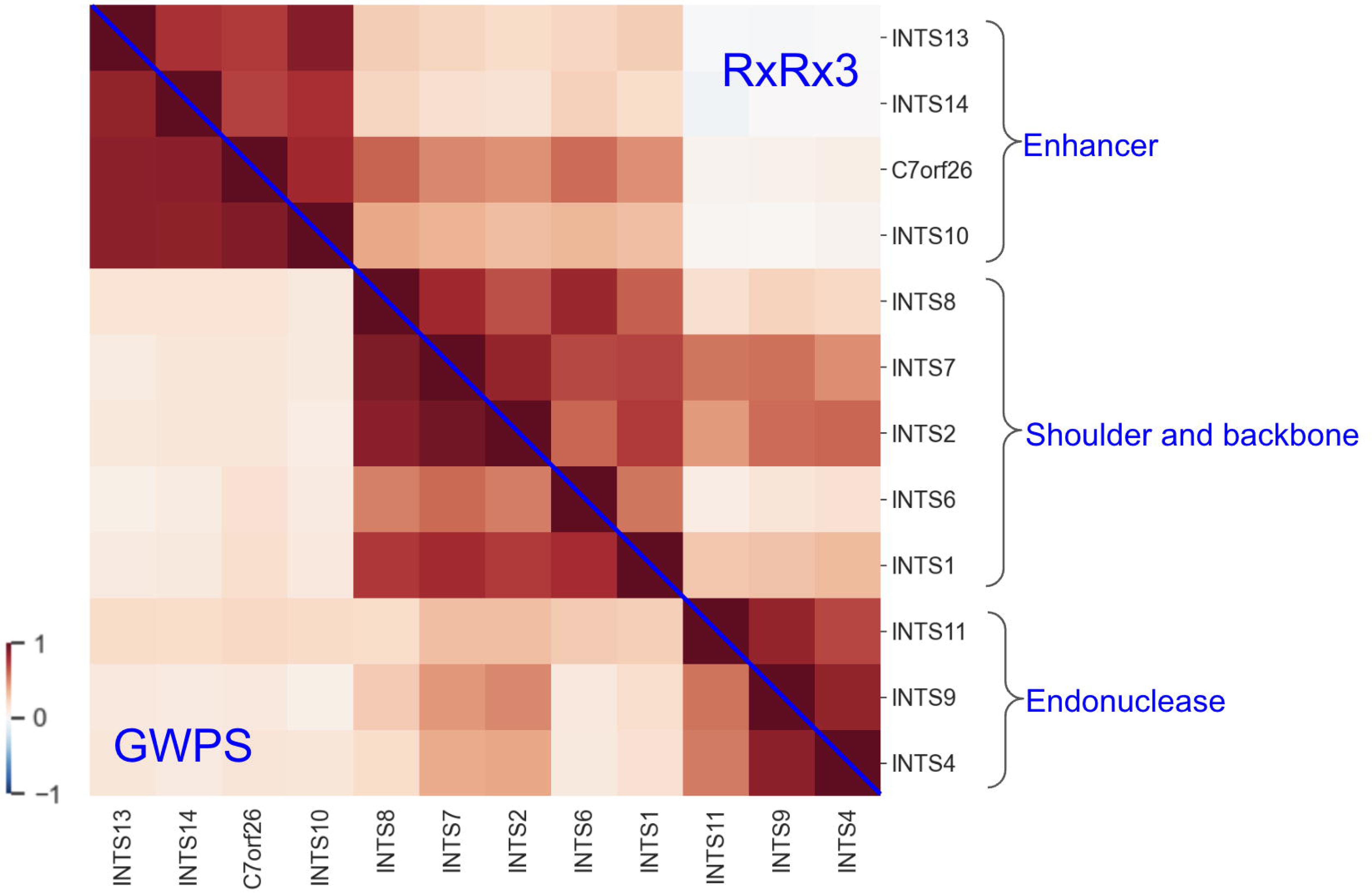
A split cosine similarity heatmap of the Integrator complex subunits from the RxRx3 and GWPS maps. Above the diagonal represents similarities for the RxRx3 map, and below the diagonal represents similarities for the GWPS map. There are three main clusters visible in both, which correspond to the three main modules of the Integrator complex.

#### 4.3.5 Evidence for the roles of uncharacterized genes *C18orf21* and *C1orf131*

Motivated by the shared cosine similarity structure of the Integrator complex in the GWPS and RxRx3 maps and a suggested role for the recently characterized *C7orf26*, we sought to determine whether there are any other uncharacterized genes whose roles might be elucidated using these maps. We focused on two uncharacterized genes: *C18orf21* and *C1orf131* and examined their top connections in the PCA-TVN maps of RxRx3 and GWPS.

Six genes are in the top 25 most cosine similar genes to *C18orf21* in both GWPS and RxRx3 maps (Figure 8a): *ABT1*, *DKC1*, *POP1*, *POP5*, *RPP30*, and *UTP23*. We performed a gene set enrichment analysis on Gene Ontology (GO) biological process, cellular component, and molecular function annotations from MSigDB ([39]) and identified terms involved in various RNA-related processes and complexes (Figure 8b). Key roles include the formation and function of ribonucle-oprotein complexes (SNO_S_RNA_CONTAINING_RIBONUCLEOPROTEIN_COMPLEX, MULTIMERIC_RIBONUCLEASE_P_COMPLEX) and critical RNA maturation steps (RNA_5_END_PROCESSING, RNA_CAPPING, tRNA_5_LEADER_REMOVAL). These genes are also crucial for ribosome biogenesis (RIBOSOME_BIOGENESIS) and exhibit endonuclease activities (RIBONUCLEASE_P_ACTIVITY). The involvement in the nucleolus further underscores their role in RNA processing and structural organization.

**Figure 8:**
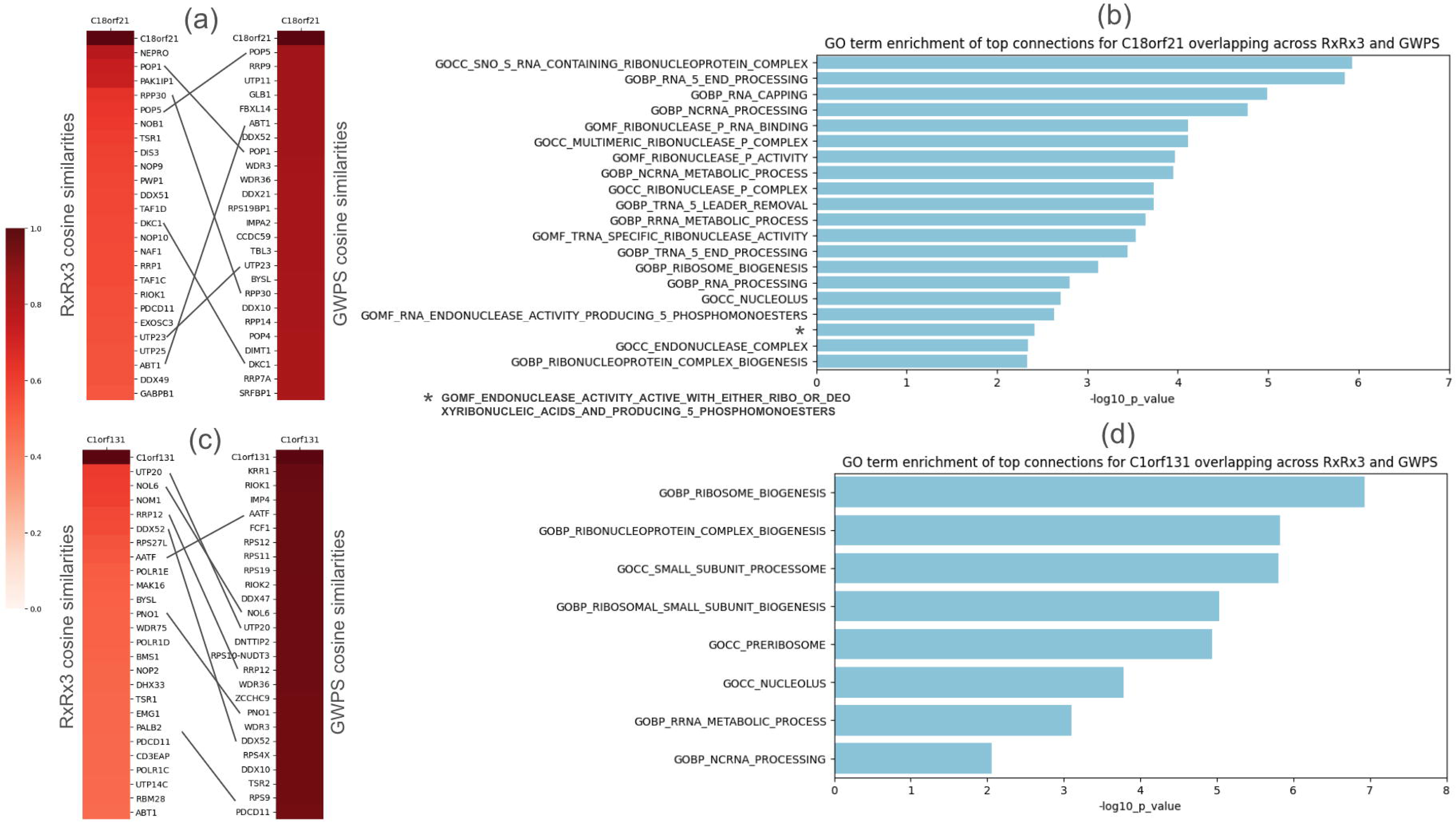
(a) Top 25 strongest connections to *C18orf21* and the associated cosine similarities in each of the RxRx3 and GWPS maps. Overlapping six genes across are connected by lines between the two heatmaps. (b) GO enrichment results for the six genes that are in the overlap of the top 25 strongest connections to *C18orf21* in the rxrx and GWPS maps. Bar lengths represent the Bonferroni-corrected -log10(p-value) from a hypergeometric test. (c-d) Similar data is presented for *C1orf131*.

The six genes connected to *C18orf21* in both RxRx3 and GWPS are also enriched for the ribonuclease mitochondrial RNA processing (RNase MRP) complex in CORUM [28] (p-value 2e-6), with three of the six genes (*POP1*, *RPP30*, and *POP5*) annotated in this complex. Recent studies have also suggested *C18orf21*’s role in the RNase MRP complex [40, 41]. Our findings reinforce these observations.

Additionally, we observed a strong overlap of interactions across the RxRx3 and GWPS maps for another uncharacterized gene, *C1orf131*. Seven genes are among the top 25 in both RxRx3 and GWPS (Figure 8c): *AATF*, *DDX52*, *NOL6*, *PDCD11*, *PNO1*, *RRP12*, and *UTP20*. The gene set enrichment analysis (Figure 8d) identified several GO terms highlighting the critical roles these genes play in ribosome biogenesis and RNA processing. Notably, terms such as GOCC_SMALL_SUBUNIT_PROCESSOME and GOBP_RIBOSOMAL_SMALL_SUBUNIT_BIOGENESIS emphasize the involvement in the formation and processing of the ribosomal small subunit (SSU) processome, a key component in ribosome assembly. Additionally, the terms GOBP_RIBOSOME_BIOGENESIS, GOBP_RIBONUCLEOPROTEIN_COMPLEX_BIOGENESIS, GOCC_PRERIBOSOME, and GOCC_NUCLEOLUS indicate participation in early ribosome assembly stages within the nucleolus. *C1orf131*, along with these seven genes, has been suggested as a structural component of the small ribosomal subunit (SSU) processome [42, 43, 1]. The evidence shared by the morphological and transcriptional maps presented here corroborate a role for *C1orf131* in the SSU processome. Although these maps suggest roles for these poorly characterized genes, further biochemical research is recommended to validate these hypotheses.

## 5 Discussion

In this work we present a general framework for systematically constructing whole-genome perturbative maps of biology and benchmarking their performance globally with publicly-available annotation datasets. As an application of this framework, we utilize several analytical choices to construct 18 maps from four perturbation datasets that use three distinct data types: single-cell transcriptomic data treated with CRISPR interference (GWPS), arrayed phenotypic screening with CRISPR-Cas9 knockout perturbations (RxRx3 and cpg0016), and pooled optical screening with CRISPR-Cas9 knockouts (cpg0021). Additionally, we provide standard benchmarks to assess each map’s perturbation magnitude, consistency, and recapitulation rate of known biology. Next, we compared which areas of biology are captured by the best-performing map for each dataset, and present evidence for the roles of two poorly annotated genes, *C18orf21* and *C1orf131*. Our results corroborate recent studies [28, 40, 41, 42, 43, 1] by demonstrating connections to the same set of well-characterized genes across powerful maps from two datasets (GWPS and RxRx3).

While we attempted to minimize computational pipeline differences across the maps through consistent EFAAR choices, these maps utilize different experimental methods, cell types, and study designs, all of which impact the benchmarks. For instance, cpg0016 and RxRx3 both involve arrayed assays, but cpg0016 incorporates multiple guide RNAs targeting the same gene in a single well, whereas RxRx3 adds a single guide per well and subsequently aggregates the guides together. Employing a single guide allows for more control and dataset clean-up, while targeting multiple guides may result in higher knockdown and more consistent replicates per gene perturbation. Additionally, the delivery of guides can be performed in a large pool (cpg0021 and GWPS) or through lab automation techniques in an arrayed format (cpg0016 and RxRx3). While pooling is generally more cost-efficient, mixing cells with different treatments may prevent or suppress effects that are visible in arrayed screens from cell-cell interactions in a larger co-localized population.

Previous work for conceptualizing a pipeline for perturbative map processing includes Pycytominer (github.com/cytomining/pycytominer) [44]. Pycytominer is open-source software aimed at standardizing the analysis of high-dimensional morphological features, and it has been utilized by several studies [2, 6, 45] in conjunction with CellProfiler [10] or DeepProfiler [46]. Pycytominer was designed specifically for image-based features and concentrates on individual perturbations rather than assessing the recapitulation of relationships at a map scale.

The framework presented here can be used for any large-scale biological map building and benchmarking effort regardless of data type and can be expanded to include settings where additional perturbation types (small molecules, proteins, antibodies, viruses, etc.) or assay variables (growth conditions, reagent timing, etc.) are assessed. Additionally, exploration of genome-wide data using these tools can reveal subtle sources of bias that would not be visible at a smaller scale. For example, a recent publication Lazar et al. [47] using the framework provided here, unveiled a proximity bias effect, where CRISPR knockouts display unexpected similarities to knockouts of genetically unrelated genes on the same chromosome arm.

While our framework can enable building, benchmarking, and exploring maps based on perturbation data from different modalities, we recognize that specific biological questions may require further analysis for validation. Perturbation experiments with transcriptomic readouts, like Perturb-seq, have the unique strength of identifying subsets of interpretable features driving relationships. However, this direction was not pursued in this paper, due to the aim of establishing a unified approach widely applicable across different readout technologies. For a deeper analysis of transcriptomic data we recommend resources such as MAST [48] and SCEPTRE [49].

Our study reveals important computational and biological insights and suggests opportunities for further exploration. Testing various methods more thoroughly at each stage of EFAAR could improve map quality. Our code repository is open for community contributions, including new map construction and evaluation methods, perturbation datasets, and benchmark sources. Going forward, as more genome-scale datasets are generated and published, utilizing and improving the EFAAR framework and codebase for consistent analyses could lead to new discoveries. As predictions from these maps are validated through orthogonal experimental techniques and annotation sets are improved, analyses using the tools presented here will enable a finer understanding of the relative strengths for different perturbation modalities and techniques. Moreover, as methods to combine different data types are developed, this framework can be used to standardize and evaluate multimodal explorations. Such methods present an exciting new frontier in understanding cellular biology at the whole-genome scale.

## 6 Materials and Methods

### 6.1 Perturbation data collection

For RxRx3, we downloaded the embeddings on six-channel images of wells from https://rxrx3.rxrx.ai/downloads. These embeddings were generated by passing images through a weakly-supervised convolutional neural network (CNN) pre-trained on a set of proprietary Recursion data, and the activation values from an intermediate layer of the network were utilized as the 128-dimensional embedding of each image. The model was trained to be partially resilient to batch effects as it attempts to place perturbations from different batches into the same class [36]. The first layer in the CNN applies channel intensity normalization.

For GWPS, we downloaded the raw pre-filtered single-cell RNA-seq count data for K562 cells from https://plus.figshare.com/articles/dataset/_Mapping_information-rich_ genotype-phenotype_landscapes_with_genome-scale_Perturb-seq_Replogle_et_al_ 2022_processed_Perturb-seq_datasets/20029387.

For cpg0016, we downloaded the metadata from https://zenodo.org/records/7661296/files/jump-cellpainting/metadata-v0.5.0.zip?download=1 and the CellProfiler features corresponding to the images from the CRISPR plates from https://cellpainting-gallery.s3.amazonaws.com/index.html#cpg0016-jump/. We filter CellProfiler features based on image intensity and cell count (Section 6.4).

For cpg0021, we downloaded guide-level normalized HeLa CellProfiler features corresponding to the images from the DMEM (Dulbecco’s Modified Eagle Medium) plates at https://cellpainting-gallery.s3.amazonaws.com/index.html#cpg0021-periscope/broad/workspace/profiles/HeLa/. These features were previously summarized to the level of guide-batch pairs, which is the input to our analysis.

### 6.2 Filtering genes based on expression

For RxRx3 and cpg0016, we utilized HUVEC and U2OS RNA-seq data generated using standard techniques. Unexpressed genes are defined as those with zFPKM ([50]) less than -3 in normalized bulk RNA-seq data. For cpg0021, we employed the same expression collection strategy as outlined by Ramezani et al. [2]. We used HeLa expression data from the Broad Institute’s Dependency Map (DepMap) [51], defining unexpressed genes as those with a TPM (transcripts per million) of zero. For GWPS, we treated all perturbed genes as expressed since Replogle et al. [1] reported perturbation of the expressed genome in this dataset.

### 6.3 scVI network architecture

The network is conditioned on the batch information (in this case, the sequencing channel) to correct for batch effects while identifying a lower-dimensional embedding space. We employed an embedding network that has one hidden layer with 256 nodes and a final latent representation with 128 nodes. The decoder network has the same number of latent and hidden nodes and does not share weights with the encoder. For the larger models reported in Supplementary Table 2, we also utilized twice as many nodes in the hidden layer as in the latent representation. “scvi-tools” package (version 1.1.2) [52] was utilized in our implementation of scVI models. We used default parameters for model training. The only parameters we configured are batch key, hidden layer size, and latent layer size.

### 6.4 Filtering CellProfiler features for cpg0016

We employed the Elliptic Envelope [53] method with a contamination rate of 1% to identify and eliminate outliers in features related to image intensity, specifically targeting columns containing ’ImageQuality_MeanIntensity’ in their names. Elliptic Envelope uses covariance estimation to fit a Gaussian distribution to the features. A fixed set of outliers are removed based on a level set of the density of the estimated distribution..

Subsequently, an outlier detection was applied on cell count, by retaining only the wells containing between 50 and 350 (inclusive) in the ’Cytoplasm_Number_Object_Number’ and ’Nuclei_Number_Object_Number’ columns. Supplementary Figure 8 shows the histogram of the values for ’Cytoplasm_Number_Object_Number’ (a) and ’Nuclei_Number_Object_Number’ (b) before this filtering step.

Lastly, image features were dropped by removing columns whose names start with ’Image_’. The features that start with “Image_” in CellProfiler refer to measurements and properties that are calculated at the whole image level, and they don’t hone in on specific objects or cells within the image. Despite the intensity variation correction implemented by the authors of the cpg0016 data before segmentation, we believe that image features could capture batch effect-related signals.

### 6.5 Benchmark data collection

For StringDB, we downloaded v12.0 protein links from https://string-db.org/cgi/download and classified links with a score ≥ 950 as the known relationships.

For (CORUM [28] and HuMAP [29]), we downloaded the gene clusters representing protein complexes from the online data supplementary to the corresponding study. The pairwise relationships used for biological relationship benchmark recall computation consist of the union of all pairwise within-complex relationships across all complexes.

For Reactome, we downloaded “Human protein-protein interactions” from “Protein-Protein Interactions derived from Reactome pathways” at https://reactome.org/download-data in February 2021.

For SIGNOR, we downloaded complete Homo sapiens data at https://signor.uniroma2.it/downloads.php in February 2021.

### 6.6 Perturbation signal consistency

We introduced the following notation. For a genetic perturbation *g*, we assume access to a total number of *n_g_* query perturbation units. For each perturbation unit *i* = 1*, . . ., n_g_*, we have an embedding vector *x_g,i_*. Moreover, each perturbation unit is associated with a batch *b_g,i_* ∈ {1*, . . ., B*}.

Let *g_b_* denote all perturbation units of *g* in batch *b*, and let |*g_b_*| = *n_g,b_*. Thus, 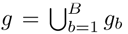 and |*g*| = *n_g_*.

As the perturbation consistency test statistic, we used avgsim, defined as the mean of the cosine similarities across all pairs of the perturbation unit profiles for *g* (i.e., in all batches). Formally,

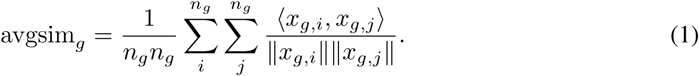

### 6.7 Perturbation signal magnitude

The energy distance [26, 27] measures how distant the replicate units of a perturbation are from the negative controls, essentially measuring the effect size of the perturbation in a high-dimensional space. For each query perturbation, we compute the distance of the replicate perturbation units’ distribution to the control units’ distribution using tests derived from energy statistics. Assuming access to two sets of embeddings *x*_1_*, . . ., x_n_*_1_ (representing query perturbation units) and *y*_1_*, . . ., y_n_*_2_ (representing negative control units), the energy distance is defined as

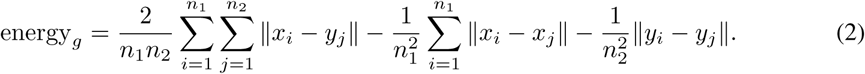

This distance will be zero when the distributions are identical, and positive between non-identical distributions.

### 6.8 Identification of significance of perturbation signal

The genes that achieved a consistency (or magnitude) p-value < .05 were considered significant. Perturbation signal benchmark reports the fraction of such genes.

Statistical significance was assessed using a permutation test comparing the test statistic (consistency or magnitude as computed above) of the query gene against an empirical null distribution generated using the corresponding metric for the set of unexpressed genes in each map. A non-parametric test was used to assign a p-value. Parametric tests are not preferred for perturbation signal benchmark metrics because the underlying population of samples do not typically follow a probability distribution from an easily-specified parametric family.

For the consistency metric, average cosine similarity was computed for the unexpressed genes *g^′^_k_*, *k* = 1*, . . ., K*. A p-value was assigned to an expressed gene *g* by

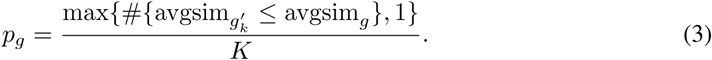

For the magnitude metric, energy distance was computed for the unexpressed genes *g^′^*, *k* = 1*, . . ., K*.

A p-value was assigned to an expressed gene *g* by

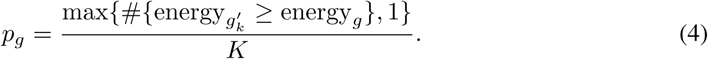

For both consistency and magnitude, at least 1000 unexpressed genes (i.e., *K* = 1000) are required for the null to get a representative p-value.

### 6.9 Recall computation

To assess how well a map recapitulates known biological relationships, we calculated recall measures on known gene-gene relationships as follows. First, we calculated pairwise cosine similarities between the aggregated perturbation embeddings of all perturbed genes in the map. Self-links were excluded since the cosine similarity for these is one and this biases the recall computation. Next, all relationships that fall below the 5th and above the 95th percentiles of this distribution were selected as “predicted links”. Then we calculated the recall for each benchmark annotation source as the ratio of the predicted true links over all links. Since different perturbation sets contain different number of genes (“Total perturbed genes” in Table 1), when we compute recall, we adjusted the denominator to only include gene-gene interactions from the benchmark annotation source in which both genes are present in the perturbation dataset. This adjustment is necessary to ensure fairness across different perturbative maps when benchmarking them against biological relationships. We then multiplied the ratios by 100 which shifts the results from ratios to percentages.

### 6.10 Noise sensitivity analyses

Subsampling was done as follows for each dataset. We repeated random selection three times and report mean and standard deviation of the results.

GWPS data contains 267 Gel Bead-in Emulsion (GEM) groups as batches. Each gene readout comes from about 250 cells. The number of GEM groups for each gene can significantly vary from two to nearly all groups. We randomly sampled 200, 100, or 50 cells for each gene across all possible GEM groups for that gene.

RxRx3 data covers ∼17,000 genes through 88 experiment templates each of which contains readouts from six CRISPR guides for each of ∼200 genes. Each experiment contains identical nine plates, and each experiment template was screened twice, resulting in a total of 176 experiments [5]. We randomly sampled two, four, or eight plate replicates for each gene from each experiment.

cpg0016 data includes six batches (runs), five of which include all of the approximately 8,000 screened genes, containing a total of about 28 plates. We randomly sampled one, two, or four out of the six runs.

cpg0021 data includes five DMEM plates, each containing readouts from four CRISPR guides for ∼20,000 genes. We randomly sampled one, two, or four out of the five plates.

### 6.11 Protein complex identification

To evaluate the accuracy of a map in recapitulating established biological gene clusters and to gain insights into its efficacy in capturing various facets of biology, we conducted the following analysis. For each known cluster with at least ten genes in the map, we generated all pairs of genes in the cluster (excluding self-links) as our ground truth for known gene relationships within that cluster. Subsequently, we computed the Kolmogorov-Smirnov (KS) statistic and its associated p-value by comparing the cosine similarity distribution of these links to the cosine similarity distribution of those that consist of one gene inside the cluster and another outside it. The latter serves as our “negative” distribution. We then defined the identified clusters as those with a p-value < .01 from each map.

### 6.12 Gene Ontology enrichment analysis

We downloaded the “GO: Gene Ontology gene sets” collection from the Molecular Signatures Database (MSigDB) under the “C5: ontology gene sets” category (https://www.gsea-msigdb.org/gsea/msigdb/collections.jsp). A hypergeometric test was performed to assess the enrichment of Gene Ontology (GO) terms within the top 25 connections of *C18orf21* or *C1orf131*. Only the terms with at least ten genes in the map were evaluated. To account for the issue of multiple hypothesis testing, Bonferroni correction was applied, ensuring a stringent control of the family-wise error rate. Only those GO terms with a corrected p-value < .01 were considered significantly enriched.

### 6.13 Code availability

The code used in this study is available for public use at the following repository: https://github.com/recursionpharma/EFAAR_benchmarking

## Acknowledgments

We would like to thank James Taylor, Renat Khaliullin, Seyhmus Guler, and John Urbanik for their contributions to the development of the EFAAR pipeline and benchmarking methodologies. We would like to thank Leslie Gaffney and Orit Rozenblatt-Rosen for their help with the graphical representation of the EFAAR pipeline. We also would like to thank Dan Maljovec for his invaluable assistance in making the benchmarking repository accessible to the public.

## Supplementary Table Legends

Supplementary Table 1: Details on cell painting protocols for RxRx3, cpg0016, and cpg0021 datasets. Although cpg0016 and RxRx3 employ the same six stains, the WGA and Phalloidin are imaged in two separate channels in RxRx3 while they are combined in a single channel in cpg0016.

Supplementary Table 2: Biological relationship benchmarks for GWPS, cpg0016, and cpg0021 datasets when using larger dimension sizes for the embedding space.

Supplementary Table 3: The complexes that are significantly identified by all three maps and the significance level for each in different maps. ***** means < 1e-5, **** means < 1e-4, *** means < 1e-3, and ** means < .01.

Supplementary Table 4: The complexes that are uniquely identified by the GWPS map and the significance level for each in different maps. ***** means < 1e-5, **** means < 1e-4, *** means < 1e-3, and ** means < .01, * means < .05, and “ns” means a non-significant p-value (≥ .05). Dashes represent the cases where the corresponding map did not have ten expressed genes from the complex, so the p-value was not calculated.

Supplementary Table 5: The complexes that are uniquely identified by the cpg0016 map and the significance level for each in different maps. ** means < .01, * means < .05, and “ns” stands for a non-significant p-value (≥ .05).

Supplementary Table 6: The complexes that are uniquely identified by the cpg0021 map and the significance level for each in different maps. ** means < .01, * means < .05, and “ns” stands for a non-significant p-value (≥ .05). Dashes represent the cases where the corresponding map did not have ten expressed genes from the complex, so the p-value was not calculated.

## Supplementary Figure Legends

Supplementary Figure 1: Visual description of (a) consistency and (b) magnitude measures.

Supplementary Figure 2: UpSet plot of the intersection of the interacting gene pairs based on the five different annotation sources we utilize in our analyses. y-axis is in log scale.

Supplementary Figure 3: Noise sensitivity analyses through down-sampling of the replicates for RxRx3. Each bar represents an average of results from three random samples. Dotted line represents the performance of a random, uninformative map.

Supplementary Figure 4: Noise sensitivity analyses through down-sampling of the replicates for GWPS. Each bar represents an average of results from three random samples. Dotted line represents the performance of a random, uninformative map.

Supplementary Figure 5: Noise sensitivity analyses through down-sampling of the replicates for cpg0016. Each bar represents an average of results from three random samples. Dotted line represents the performance of a random, uninformative map.

Supplementary Figure 6: Noise sensitivity analyses through down-sampling of the replicates for cpg0021. Each bar represents an average of results from three random samples. Dotted line represents the performance of a random, uninformative map.

Supplementary Figure 7: Integrator complex heatmap for the cpg0021 PCA-TVN map. The modular structure of the Integrator complex is not visible.

Supplementary Figure 8: (a) Cytoplasm and (b) nuclei object counts for the cpg0016 data.

## Notes

### Competing Interest Statement

All authors are current or former employees of Recursion Pharmaceuticals, Inc. or Genentech, Inc., and have received real or optional ownership interest in the companies.

### Summary of Updates

- We added more datasets and more EFAAR pipelines using which we build and benchmark perturbative maps. - We added additional analyses to explore the built maps to discover new biology. - We clarified the goals and contributions of the work.

