## Supplemental for "Building, Benchmarking, and Exploring Perturbative Maps of Transcriptional and Morphological Data"

Supplementary Table 1

| | Channel count | Stains | Cellular compartments labeled |
|---|---|---|---|
| RxRx3 | 6 | MitoTracker<br>Hoechst<br>SYTO 14<br>Concanavalin A<br>Wheat germ agglutinin (WGA)<br>Phalloidin | Mitochondria<br>Nucleus<br>Nucleoli and cytoplasmic RNA<br>Endoplasmic reticulum<br>Golgi and cell membrane<br>Actin filaments |
| cpg0016 | 5 | MitoTracker<br>Hoechst<br>SYTO 14<br>Concanavalin A<br>Wheat germ agglutinin (WGA)<br>Phalloidin | Mitochondria<br>Nucleus<br>Nucleoli and cytoplasmic RNA<br>Endoplasmic reticulum<br>Golgi and cell membrane<br>Actin filaments |
| cpg0021 | 5 | Anti-TOMM20 108 antibody<br>DAPI<br>Concanavalin A<br>Wheat germ agglutinin (WGA)<br>Phalloidin | Mitochondria<br>Nucleus<br>Endoplasmic reticulum<br>Golgi and cell membrane<br>Actin filaments |

Supplementary Table 2

(a) GWPS

| | | CORUM | HuMAP | Reactome | SIGNOR | StringDB |
|---|---|---|---|---|---|---|
| 128D | scVI | 14.7% | 21.6% | 7.7% | 7.3% | 18.2% |
| | scVI-CS | 47.3% | 36.8% | 21.1% | 13.3% | 48.7% |
| | scVI-TVN | 47.4% | 36.8% | 21.8% | 13.7% | 49.2% |
| | PCA | 26.0% | 20.7% | 13.3% | 13.3% | 29.8% |
| | PCA-CS | 48.1% | 36.3% | 21.9% | 14.0% | 49.0% |
| | PCA-TVN | 49.1% | 38.2% | 21.6% | 13.7% | 49.9% |
| 256D | scVI | 15.1% | 20.6% | 8.3% | 7.4% | 17.4% |
| | scVI-CS | 48.2% | 37.1% | 22.6% | 14.7% | 49.7% |
| | scVI-TVN | 48.1% | 37.5% | 21.6% | 13.7% | 49.6% |
| | PCA | 27.2% | 21.6% | 13.6% | 13.8% | 30.9% |
| | PCA-CS | 48.8% | 37.0% | 22.4% | 14.0% | 49.4% |
| | PCA-TVN | 49.7% | 38.8% | 21.9% | 14.0% | 50.3% |
| 512D | scVI | 15.1% | 18.4% | 8.9% | 7.4% | 16.0% |
| | scVI-CS | 49.2% | 38.7% | 23.1% | 14.4% | 50.3% |
| | scVI-TVN | 48.5% | 37.9% | 22.4% | 13.8% | 49.7% |
| | PCA | 28.7% | 22.9% | 14.3% | 14.0% | 32.3% |
| | PCA-CS | 49.4% | 37.5% | 21.4% | 14.2% | 49.6% |
| | PCA-TVN | 50.2% | 39.6% | 21.4% | 13.8% | 50.5% |

(b) cpg0016

| | | CORUM | HuMAP | Reactome | SIGNOR | StringDB |
|---|---|---|---|---|---|---|
| 128D | PCA | 12.7% | 14.2% | 10.4% | 9.8% | 13.1% |
| | PCA-CS | 21.3% | 17.8% | 14.1% | 12.3% | 19.1% |
| | PCA-TVN | 22.7% | 19.2% | 14.6% | 12.8% | 19.8% |
| 256D | PCA | 12.7% | 14.2% | 10.5% | 9.7% | 13.2% |
| | PCA-CS | 22.5% | 18.9% | 14.9% | 12.6% | 20.3% |
| | PCA-TVN | 24.2% | 20.1% | 15.5% | 12.9% | 20.7% |
| 512D | PCA | 12.8% | 14.2% | 10.6% | 9.7% | 13.3% |
| | PCA-CS | 23.2% | 19.5% | 15.9% | 13.4% | 21.3% |
| | PCA-TVN | 26.0% | 21.2% | 16.3% | 13.3% | 22.0% |

(c) cpg0021

| | | CORUM | HuMAP | Reactome | SIGNOR | StringDB |
|---|---|---|---|---|---|---|
| 128D | PCA | 17.4% | 15.0% | 12.4% | 12.1% | 18.8% |
| | PCA-CS | 19.4% | 16.2% | 13.9% | 12.6% | 19.6% |
| | PCA-TVN | 24.7% | 15.5% | 14.9% | 16.7% | 17.1% |
| 256D | PCA | 17.9% | 15.4% | 12.8% | 12.3% | 19.3% |
| | PCA-CS | 18.8% | 15.8% | 13.5% | 12.0% | 19.4% |
| | PCA-TVN | 25.2% | 16.1% | 16.3% | 17.6% | 18.1% |
| 512D | PCA | 18.2% | 15.7% | 12.9% | 12.4% | 19.6% |
| | PCA-CS | 18.5% | 15.9% | 13.1% | 11.5% | 19.4% |
| | PCA-TVN | 19.0% | 13.3% | 13.8% | 15.2% | 14.5% |

Supplementary Table 3

| Protein Complex | GWPS | cpg0016 | cpg0021 |
|---|---|---|---|
| 20S proteasome | ***** | ***** | ** |
| BRCA1-core RNA polymerase II complex | ***** | ***** | **** |
| C complex spliceosome | ***** | ***** | ***** |
| DNA synthesome complex (13 subunits) | ***** | ***** | ** |
| Mediator complex | ***** | ***** | ** |
| Nop56p-associated pre-rRNA complex | ***** | **** | ***** |
| PA28-20S proteasome | ***** | ***** | ** |
| PA700-20S-PA28 complex | ***** | ***** | **** |
| RNA polymerase II core complex | ***** | ***** | *** |
| Respiratory chain complex I (beta subunit) mitochondrial | ***** | *** | ** |
| Respiratory chain complex I (holoenzyme), mitochondrial | ***** | ***** | ***** |
| Spliceosome | ***** | ***** | ***** |

Supplementary Table 4

| Protein Complex | GWPS | cpg0016 | cpg0021 |
|---|---|---|---|
| TFIIH transcription factor complex | ***** | - | ns |
| Exosome | ***** | - | ns |
| CRSP-Mediator 2 complex | ***** | - | ns |
| PC2 complex | ***** | - | ns |
| TRAP-SMCC mediator complex | ***** | - | ns |
| Integrator-RNAPII complex | ***** | - | ns |
| Integrator complex | ***** | - | ns |
| CSA complex | ***** | - | ns |
| DDB2 complex | ***** | - | ns |
| DSS1 complex | ***** | - | ns |
| CSA-POLIIa complex | ***** | - | ns |
| STAGA complex | ***** | - | ns |
| 12S U11 snRNP | ***** | - | ns |
| INO80 chromatin remodeling complex | ***** | - | ns |
| Cytochrome c oxidase, mitochondrial | ***** | - | ns |
| SWI-SNF chromatin remodeling-related-BRCA1 complex | ***** | ns | ns |
| CENP-A nucleosomal complex | ***** | - | * |
| PBAF complex (Polybromo- and BAF containing complex) | ***** | - | ns |
| BAF complex | ***** | ns | ns |
| TNF-alpha/NF-kappa B signaling complex 6 | ***** | - | * |
| EBAFa complex | ***** | ns | ns |
| EBAFb complex | ***** | ns | ns |
| BRG1-SIN3A-HDAC containing SWI/SNF remodeling complex | ***** | ns | ns |
| DMAP1-associated complex | ***** | - | ns |
| PCAF complex | ***** | - | ns |
| BRG1-SIN3A complex | ***** | ns | ns |
| BRM-SIN3A-HDAC complex | ***** | ns | ns |
| LARC complex (LCR-associated remodeling complex) | ***** | ns | ns |
| Exon junction complex | ***** | - | ns |
| CF IIAm complex (Cleavage factor IIAm complex) | ***** | - | ns |
| ING2 complex | **** | - | ns |
| mRNA decay complex (UPF1, UPF2, UPF3B, DCP2, XRN1) | **** | - | ns |
| TRAPP complex | **** | - | ns |
| ALL-1 supercomplex | **** | ns | * |
| BRM-associated complex | **** | ns | ns |
| HRD1 complex | **** | - | ns |
| ATAC complex, YEATS2-linked | *** | - | ns |
| HCF-1 complex | *** | ns | ns |
| SIN3-ING1b complex II | *** | ns | ns |
| BASC complex (BRCA1-associated genome surveillance complex) | *** | ns | ns |
| BHC110 complex | *** | - | ns |
| WINAC complex | *** | ns | ns |
| SNF2h-cohesin-NuRD complex | *** | ns | ns |
| GCN5-TRRAP histone acetyltransferase complex | ** | - | ns |
| BRM-SIN3A complex | ** | ns | * |
| v-ATPase-Ragulator-AXIN/LKB1-AMPK complex | ** | - | ns |
| Set1B complex | ** | - | ns |
| NUMAC complex (nucleosomal methylation activator complex) | ** | - | ns |
| CENP-A NAC-CAD complex | ** | - | ns |
| anti-BHC110 complex | ** | - | ns |

Supplementary Table 5

| Protein Complex | GWPS | cpg0016 | cpg0021 |
|---|---|---|---|
| CtBP complex | ns | ** | * |

Supplementary Table 6

| Protein Complex | GWPS | cpg0016 | cpg0021 |
|---|---|---|---|
| MLL1-WDR5 complex | ns | ns | ** |
| Emerin complex 25 | ns | - | ** |
| Intraflagellar transport complex B | ns | - | ** |
| AFF4 super elongation complex (SEC) | ns | * | ** |

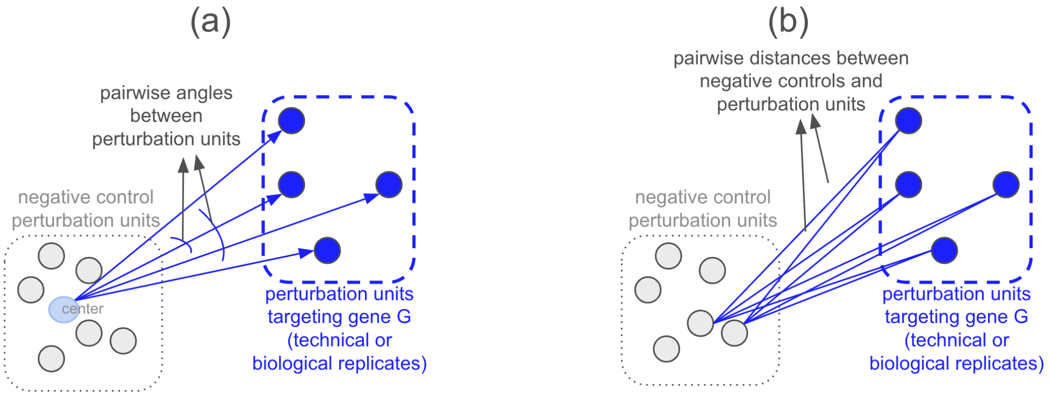


Supplementary Figure 1

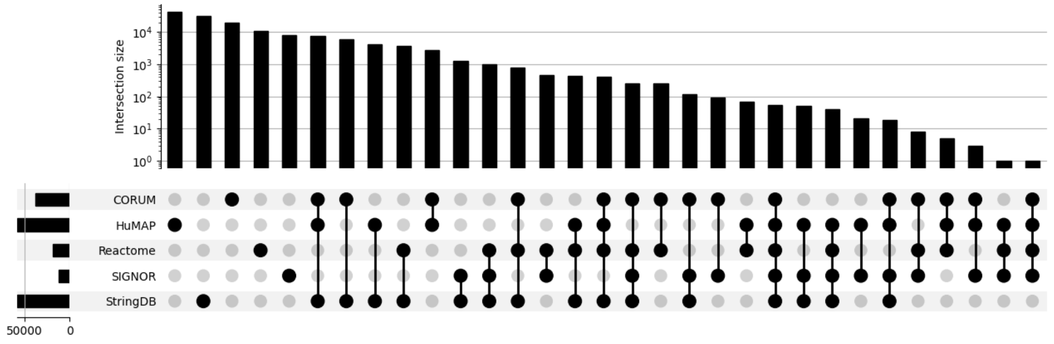


Supplementary Figure 2

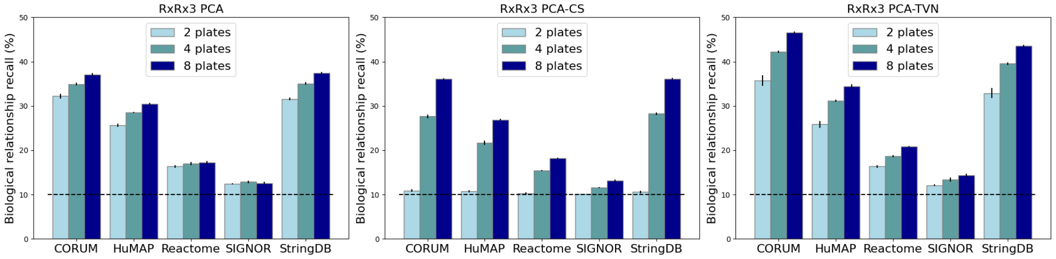


Supplementary Figure 3

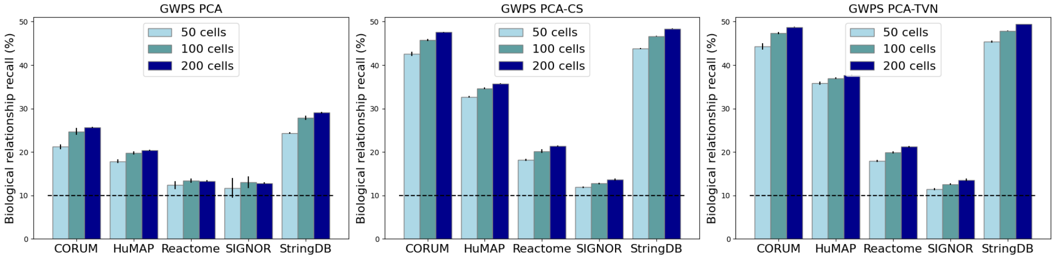


Supplementary Figure 4

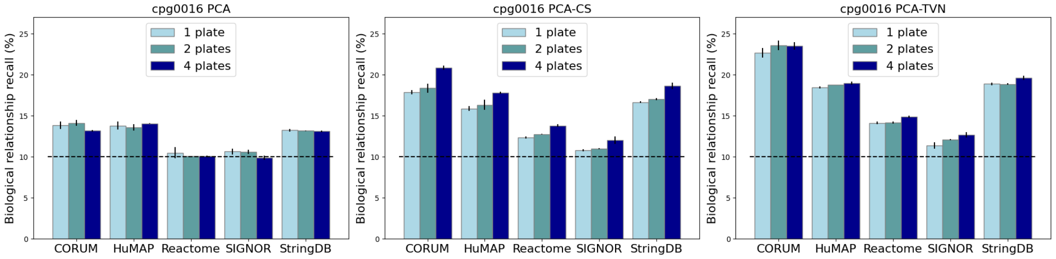


Supplementary Figure 5

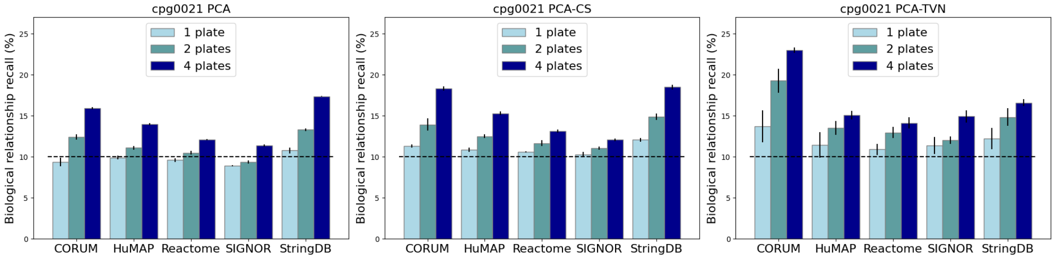


Supplementary Figure 6

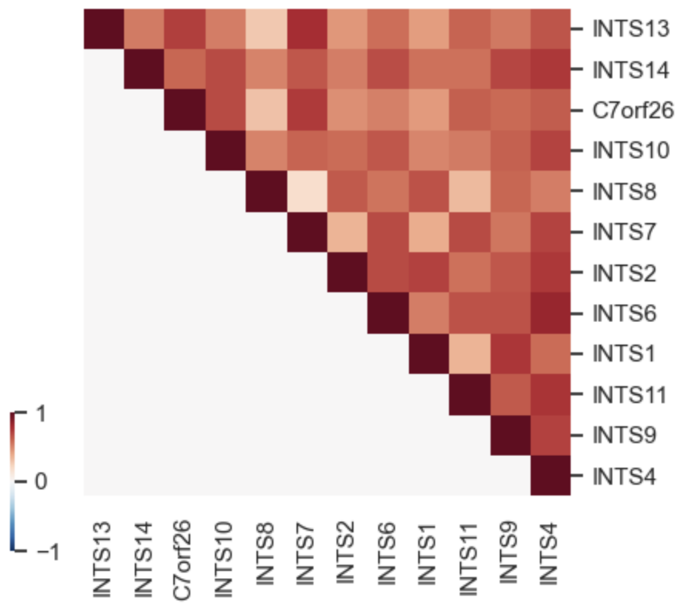


Supplementary Figure 7

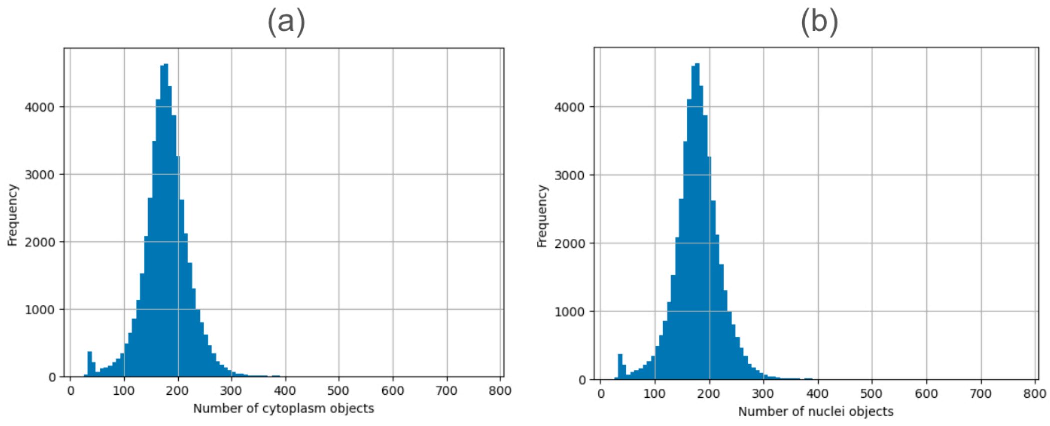


Supplementary Figure 8
